# Surface-attached model lipid membranes derived from human red blood cells

**DOI:** 10.1101/2025.08.18.670922

**Authors:** Sanyukta Prakash Mudakannavar, Matthew D. Mitchell, Robert J. Rawle

**Author notes:** Co-first author contribution (Sanyukta Prakash Mudakannavar, Matthew D. Mitchell).

## Abstract

Red blood cells (RBCs) are the most abundant human cell type and interface interactions with the RBC membrane are at the heart of many processes relevant for human health, such as immune system modulation, interactions with foreign pathogens and with pharmacological drugs. To better study such membrane interface interactions, it would be useful to employ surface - attached model lipid membranes derived from RBCs to enable surface-sensitive biophysical and biochemical measurements. Here, we present approaches to prepare two such types of RBC-derived model lipid membranes – supported lipid bilayers (RBC-SLBs) and tethered RBC liposomes. We present data characterizing and validating these model membranes, including assessing lipid mobility, the distribution and mobility of the glycophorin A membrane protein, the functionality of the acetylcholinesterase enzyme, and the utility of the RBC-SLBs as binding targets for viral pathogens. We anticipate that our results and methodologies will be of interest to researchers studying molecular interactions with RBC membranes, as well as those interested in the engineering of model membrane platforms derived from other physiological membranes.

## Introduction

Red blood cells (RBCs), or erythrocytes, are the most abundant cell type in the human body and as such are important interaction partners in a variety of processes relevant for human health, including interactions with immune system components, interactions with bacterial and viral pathogens, and interactions with small molecules, including pharmacological drugs ^1–6^. To better understand and quantify such interactions, it would be useful to employ surface-attached model lipid membranes derived from RBCs as target membranes. Such model membranes would enable precise membrane-interaction measurements by a variety of surface-sensitive techniques, including fluorescence microscopy, surface plasmon resonance, quartz crystal microbalance with dissipation monitoring, bio-layer interferometry, etc. They would also allow the researcher to exert control over membrane geometry, morphology, and external environment to facilitate measurements of biophysical and biochemical membrane properties.

Historically, surface-attached model membranes have been formed solely from purified lipids ^7,8^. Recently, however, researchers have begun developing methods to form model membranes from more complex physiologically-derived membranes ^9–12^, such as outer membrane vesicles of bacteria, isolated endosomes, and plasma membrane vesicles from mammalian cells. Although many different types of model membrane platforms exist, the two platforms which we focus on in this report are: 1) glass supported lipid bilayers (SLBs) and 2) surface-tethered liposomes. To our knowledge, neither of these model membrane platforms has been developed for RBCs.

SLBs are contiguous planar membranes formed on a glass surface, typically by self-assembly using one of many reported methodologies ^8^. An important feature of SLBs is that they preserve the lateral fluidity of the lipids in the membrane.

Surface-tethered liposomes are liposomes that have been specifically bound to a solid support, often a microscope coverslip which has been modified with a polymer or other passivating layer to prevent non-specific adherence of the liposome to the glass surface itself. A variety of attachment chemistries have been employed ^11,13–16^, such as programmable hybridization of lipid-anchored DNA oligos and lipid-anchored biotin-avidin interactions. Often, the liposomes are large unilamellar vesicles ∼100-200 nm in diameter, but giant unilamellar vesicles several µm in diameter can also be used. The goal with surface-tethered liposomes is typically to reduce the interaction with the underlying substrate compared to a supported lipid bilayer, in which close proximity to the solid support can sometimes produce unwanted interactions.

Here, we report the development of SLBs and surface-tethered liposomes derived from RBCs. We characterize each membrane type, including assessing the mobility and accessibility of membrane components. We validate the integrity/activity of key RBC components, including transmembrane proteins and membrane-anchored enzymes. We also examine binding by viral pathogens. We primarily focus on studying these model membranes via fluorescence microscopy, but the membrane preparation methods we describe could be easily employed for study by a variety of other surface-sensitive techniques, including SPR, QCM, etc.

We anticipate that this report will be useful to researchers studying interactions with RBC membranes, as well as those interested in the engineering of model membrane platforms derived from other physiological membranes.

## Results and Discussion

### Preparation of labeled RBC liposomes

In this report, we discuss the preparation of two distinct model membrane platforms derived from human red blood cells – 1) supported lipid bilayers (RBC-SLBs) and 2) tethered RBC liposomes. As a first step in the preparation of both membrane platforms, RBCs were transformed into labeled RBC liposomes (see **Figure 1** and full details in the Materials and Methods). To do this, RBCs were treated with a series of hypotonic solutions followed by purification by centrifugation. This removed the internal hemoglobin and other RBC contents, replacing them with external buffer. This process produces so-called “RBC ghosts” as previously described by a number of reports ^17–20^. As needed, the membranes of these RBC ghosts were then labeled with a lipophilic fluorescent dye (typically Oregon Green-DHPE) and/or a biotinylated lipid via incubation and passive membrane partitioning. The labeled ghosts were then transformed into labeled RBC liposomes by extrusion. In our typical preparations, these RBC liposomes were successively extruded at 400 nm and then 100 nm normal pore size, yielding a monodisperse, but somewhat broad, distribution of liposomes with an average diameter of ∼200 nm, as determined by dynamic light scattering (DLS) (**Figure S1**). These labeled RBC liposomes were then used to prepare either RBC-SLBs or tethered RBC liposomes, as described below.

**Figure 1.**
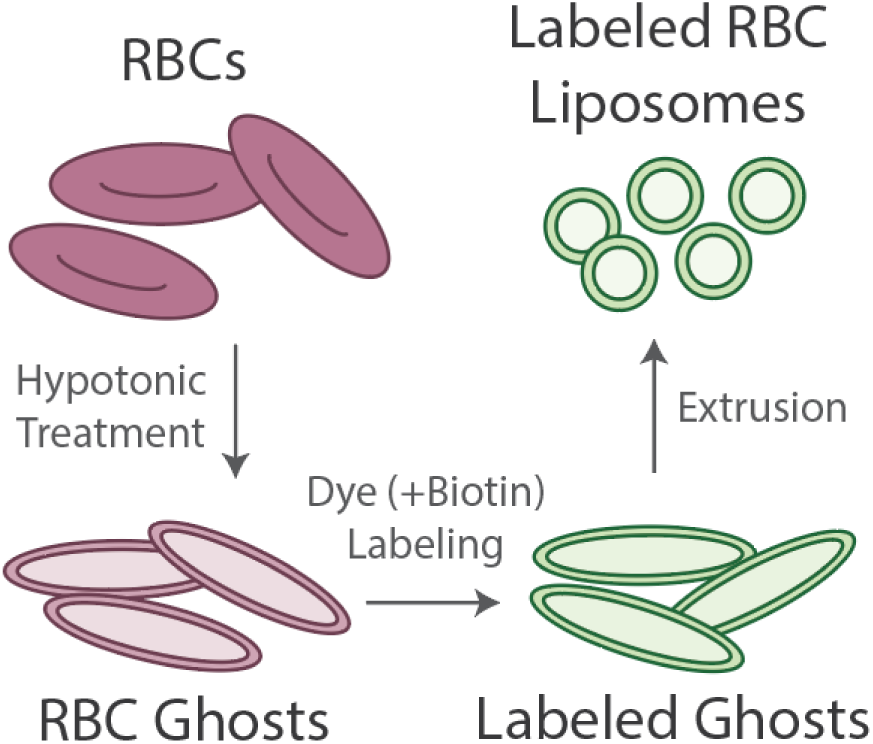
Preparation of labeled RBC liposomes. Red blood cells (RBCs) undergo treatment with a hypotonic solution and purification by centrifugation to remove hemoglobin and other internal contents. This produces RBC ghosts. Ghosts are incubated with lipophilic fluorescent dye and/or biotin-lipid, followed by purification by centrifugation, to produce labeled ghosts. Labeled ghosts then undergo extrusion to produce labeled RBC liposomes. These liposomes are then used as the starting material for the model membranes described below.

### Preparation of RBC-SLBs

We formed SLBs by the commonly-used vesicle fusion technique ^21^, in which liposomes bind to a clean glass surface, rupture, and then merge with adjacent liposomes to form a contiguous bilayer that preserves the lateral mobility of the lipid components. Our early attempts to prepare SLBs solely from labeled RBC liposomes were not successful. We observed that RBC liposomes could bind successfully to the glass coverslip, but they did not appear to rupture and/or merge effectively. This was verified quantitatively by fluorescence recovery after photobleaching (FRAP), demonstrating almost complete lack of lipid mobility (**Figure S2**).

Therefore, we turned to a strategy recently reported by Daniel and coworkers to prepare SLBs from a variety of more complex physiological membranes, including bacterial outer membrane vesicles and plasma membrane vesicles (PMVs) ^9,10^. This strategy involves utilizing “rupture vesicles” (liposomes containing a small mole percent of PEGylated lipids) to facilitate rupture and merger of the physiologically derived liposomes. It also serves to distance the resulting SLB from the underlying support due to the size of the PEG headgroup.

In our implementation of this strategy (**Figure 2**), we first bound labeled RBC liposomes to a glass coverslip inside a microfluidic flow cell. Unbound RBC liposomes were removed by rinsing, and then unlabeled rupture vesicles were introduced into the flow cell and incubated to form the SLB. This strategy proved fruitful; in our test experiments we were able to qualitatively observe recovery following photobleaching, indicating SLB formation (see **Figure 3A-C**). However, some punctate spots were still observed in the resulting SLB, indicating incomplete rupture/merger of at least some RBC liposomes.

**Figure 2.**
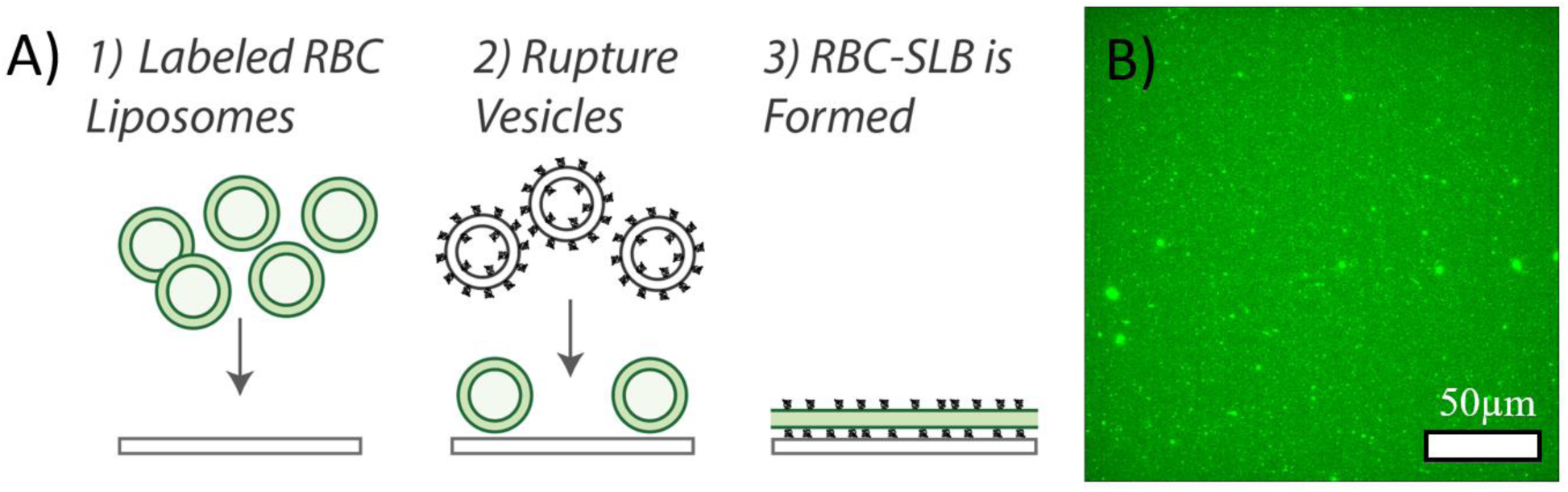
Preparation of RBC-SLBs. A) Schematic of preparation of RBC-SLBs. Labeled RBC liposomes are added to a clean glass coverslip inside a microfluidic device. Unbound liposomes are removed by buffer rinse. Rupture vesicles containing a small mole percent of PEGylated lipids are then added. These bind to empty locations on the coverslip and induce rupture/merger of neighboring RBC liposomes, resulting in the RBC-SLB. B) Example fluorescence micrograph of an RBC-SLB.

**Figure 3.**
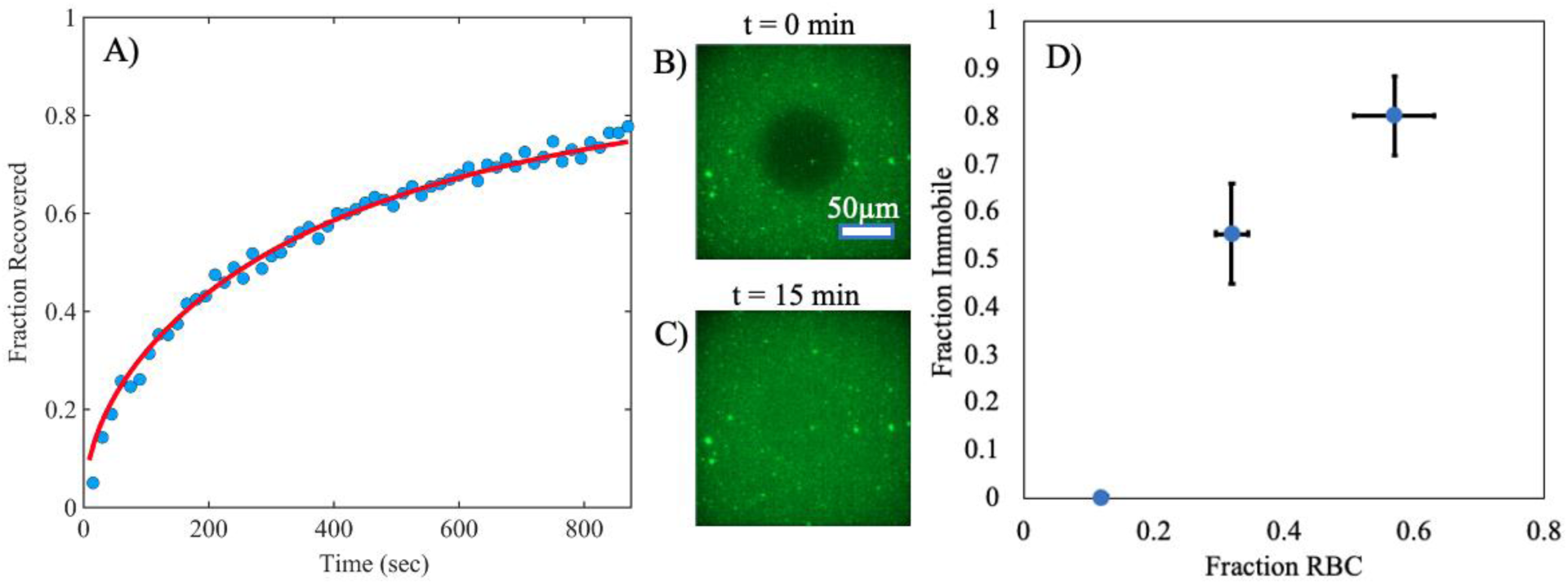
RBC-SLB mobility varies with the fraction of SLB composed of the RBC liposomes. RBC-SLBs were prepared using the rupture vesicle strategy, but with varying concentrations of added labeled liposomes. Mobility of the resulting SLB was assessed by FRAP. A) Example FRAP recovery curve (blue circles = data, red line = fit to FRAP diffusion model, **Equation 2**). Fraction recovered is the normalized fluorescence intensity within the photobleached spot, background corrected for residual photobleaching that occurred during the time-lapse imaging. Fraction recovered = 1 was set to the fluorescence intensity immediately prior to photobleaching. Fraction recovered = 0 was set to the fluorescence intensity immediately after photobleaching (t = 0). B) and C) show example fluorescence micrographs at t = 0 and t = 15 min, respectively. D) shows the dependence of the fraction immobile on the fraction of SLB composed of RBC liposomes (Fraction RBC). Fraction immobile was determined by FRAP diffusion model fits (values shown are average ± standard deviation of 3 sample replicates). Fraction RBC was determined by total image fluorescence comparisons to a standard SLB composed only of rupture vesicles (see main text and Materials and Methods for details). Values shown are average ± propagated error of standard deviations of experimental and standard SLB samples. Standard deviations were calculated from ≥ 20 image locations across 2 sample replicates.

Therefore, to quantitatively test the effect of the surface density of the bound RBC liposomes on the quality of the SLB, we prepared RBC-SLBs using the rupture vesicle strategy, but added varying concentrations of the labeled RBC liposomes (ranging from ∼0.5-5 nM). We then used FRAP to measure the mobility of the Oregon Green-DHPE lipid (which had only been included in the RBC liposomes) in the resulting RBC-SLBs.

Separately, we also quantified the fraction of the resulting RBC-SLB that originated from RBC liposomes as compared to the rupture vesicles. In this measurement, Oregon Green-DHPE was included in the rupture vesicles, whereas the RBC liposomes were left unlabeled. RBC-SLBs were prepared, and the average fluorescence within a field-of-view was quantified (*F_RBC-SLB_*). This was compared to the average fluorescence observed in an SLB composed solely of the labeled rupture vesicles (*F_rupture only SLB_*). From this, the fraction of the RBC-SLB derived from RBC liposomes could be estimated as *Fraction_RBC_*= 1 - *F_RBC-SLB_*/*F_rupture only SLB_*.

Combining both sets of data (lipid mobility and *Fraction_RBC_*), we observed a clear dependence of lipid mobility on *Fraction_RBC_*(**Figure 3D**). At low *Fraction_RBC_* ∼0.1, the immobile lipid fraction determined by FRAP was near zero. But by *Fraction_RBC_* ∼0.6, nearly all lipids were immobile (immobile fraction = 0.8 ± 0.08), suggesting mostly incomplete SLB formation. These results matched our qualitative observations of the SLB images – the density of observed punctate spots (presumably un-merged RBC liposomes) clearly increased with *Fraction_RBC_*. These spots likely account for some portion of the immobile fraction. Nonetheless, these results indicate A) that SLB formation can be achieved, but B) that the quality of the resulting SLB is dependent on the fraction of the SLB that was derived from the RBC liposomes.

There was not a strong dependence of the estimated lipid diffusion coefficient (D) on *Fraction_RBC_*; average D values for SLBs were ∼1.2-2.5 µm^2^/sec (**Figure S3**), comparable with reported measurements of lipid mobility in SLBs ^22–25^.

### Characterization/validation of RBC-SLBs

In addition to the lipid FRAP measurements, we conducted a number of experiments to validate and characterize the RBC-SLBs. Each is described separately below. These experiments were conducted using SLBs with medium *Fraction_RBC_* ∼0.3-0.4.

To characterize the distribution and accessibility of RBC membrane proteins in the resulting SLBs, we employed immunofluorescence (IF) to observe glycophorin A (GpA) – an abundant transmembrane glycoprotein in RBCs ^26^. We observed widespread immunofluorescence across the RBC-SLB, with >20x higher immunofluorescence signal compared to a control SLB composed solely of rupture vesicles (**Figure 4A**). This indicates that glycophorin A is well-distributed and accessible across the RBC-SLB. We did observe some clustered spots of immunofluorescence signal in the RBC-SLBs (see example images in **Figure 4C**), indicative of unruptured RBC liposomes and/or clustering of the GpA proteins.

**Figure 4.**
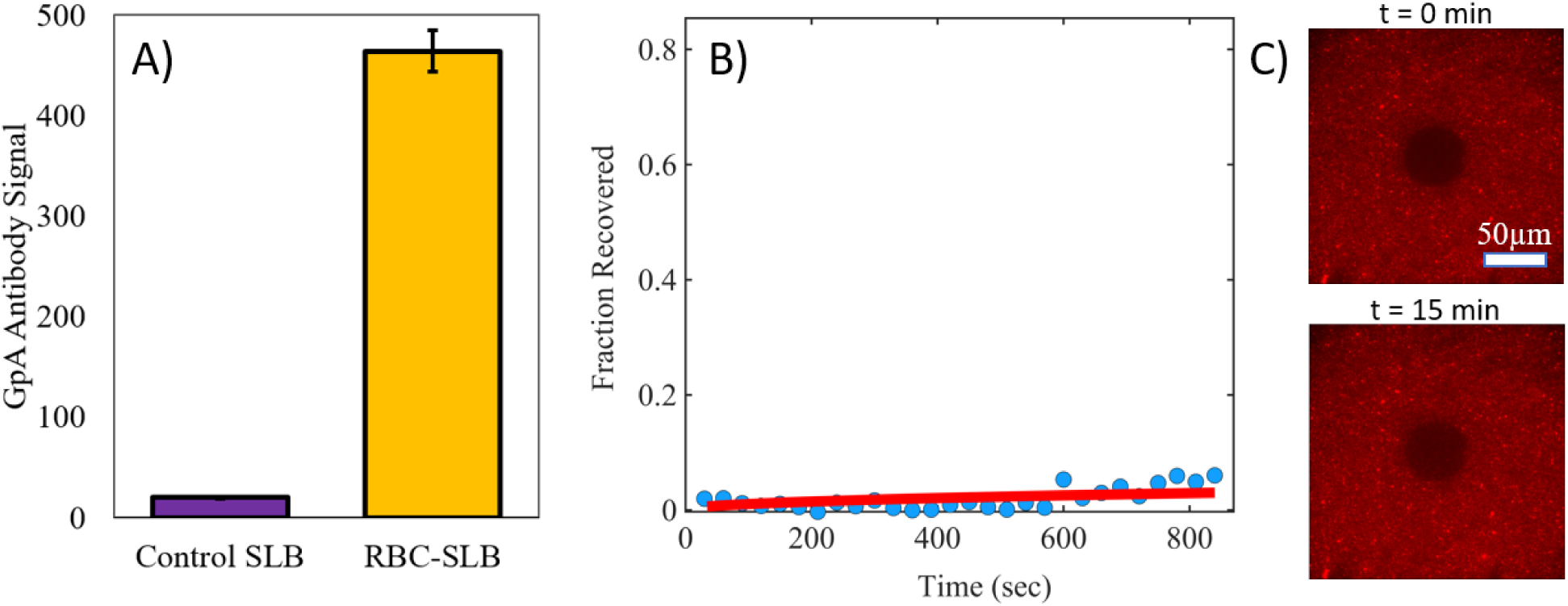
Glycophorin A in RBC-SLBs is accessible and well-distributed, but exhibits low mobility. RBC-SLBs formed using the rupture vesicle approach were immunofluorescently labeled for the glycophorin A transmembrane protein (Primary antibody = mouse IgG anti - glycophorin A (CD235a), secondary antibody = goat anti-mouse IgG with Alexa 647). A) Glycophorin A (GpA) antibody binding to RBC-SLBs or SLBs composed of rupture vesicles only (control SLB). GpA antibody signal is the Alexa 647 fluorescence intensity (average ± standard deviation of 3 sample replicates, with ≥3 image locations in each sample). B) Example FRAP recovery curve (blue circles = data, red line = fit to FRAP diffusion model, **Equation 2**). Fraction recovered is the normalized fluorescence intensity within the photobleached spot, background corrected for residual photobleaching that occurred during the time-lapse imaging. Fraction recovered = 1 was set to the fluorescence intensity immediately prior to photobleaching. Fraction recovered = 0 was set to the fluorescence intensity immediately after photobleaching (t = 0). C) Example fluorescence micrographs at t = 0 and t = 13 min, respectively.

To assess the mobility of RBC membrane proteins in the SLB, we conducted FRAP measurements on the immunolabeled glycophorin A, monitoring the recovery of the photobleached secondary antibody (**Figure 4B-C**). We observed little recovery across 15+ minutes, indicating that glycophorin A is largely immobile in the RBC-SLBs. This immobility may be due to interactions with the underlying substrate or steric corralling by other membrane components, including other membrane proteins and the PEG polymer included in the rupture vesicles. We also cannot rule out that antibody cross-linking might lead to apparent immobility.

To assess the functionality of the RBC-SLBs, we conducted two experiments. First, we quantified the enzymatic activity of acetylcholinesterase (AchE) (**Figure 5)**. AchE is a glycophosphoinositol (GPI)-anchored enzyme on the RBC surface ^27^, responsible for enzymatically cleaving the neurotransmitter acetylcholine. We utilized a commercial acetylcholinesterase activity kit which couples the cleavage of acetylcholine to the production of a fluorogenic product (AbRed), and we adapted this assay for use in our microfluidic flow cells in which the RBC-SLBs were prepared (see Materials and Methods). Using fluorescence microscopy, we compared the average AbRed fluorescence signal produced by RBC-SLBs to the fluorescence signal produced under identical conditions from an SLB composed of rupture vesicles only (**Figure 5**). We observed >20x increase of fluorescent product from the RBC-SLB compared to the control SLB, indicating robust activity of the acetylcholinesterase enzyme in the RBC-SLBs.

**Figure 5.**
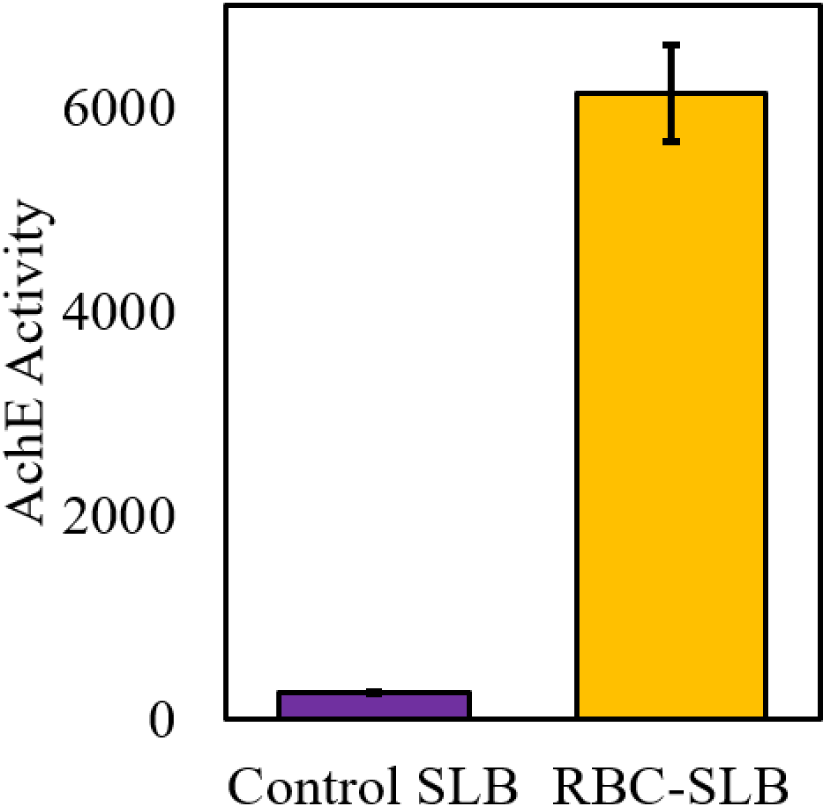
Acetylcholinesterase (AchE) activity of RBC-SLBs demonstrates that membrane-anchored enzymes are active following SLB formation. The enzymatic activity of RBC-SLBs formed using the rupture vesicle approach was assessed by a microscope-adapted red fluorogenic AchE assay (see Materials and Methods for details). This was compared to the activity of a control SLB composed of rupture vesicles only. AchE Activity is the average red fluorescence within a microscope field-of-view near the SLB surface after 5 minutes at RT. Values shown are average ± standard deviation of 3 sample replicates, with ≥10 image locations in each sample.

We note that the enzymatic activity measurement above does not rule out the possibility that some fraction of the acetylcholinesterase enzyme may be inactivated during preparation of the RBC liposomes and/or RBC-SLB. Unfortunately, there is no clear (to us) positive control SLB in which we could guarantee 100% AchE activity. However, it is possible to assess loss in enzymatic activity in the preparation of the precursor RBC liposomes due to the extrusion step, a likely candidate for activity loss should any occur. To assess this, we assayed AchE activity and specific activity of labeled RBC ghosts immediately prior to extrusion and labeled RBC liposomes immediately following extrusion (**Table 1**). We observed that the relative activity levels of the resulting RBC liposomes were comparable to the parent RBC ghosts, indicating that no substantial loss of enzymatic activity occurred during extrusion. Interestingly, the relative specific activity of the labeled RBC liposomes was substantially higher than the parent RBC ghosts, likely indicating the loss of residual hemoglobin or other non-membrane-bound protein during extrusion.

**Table 1.**
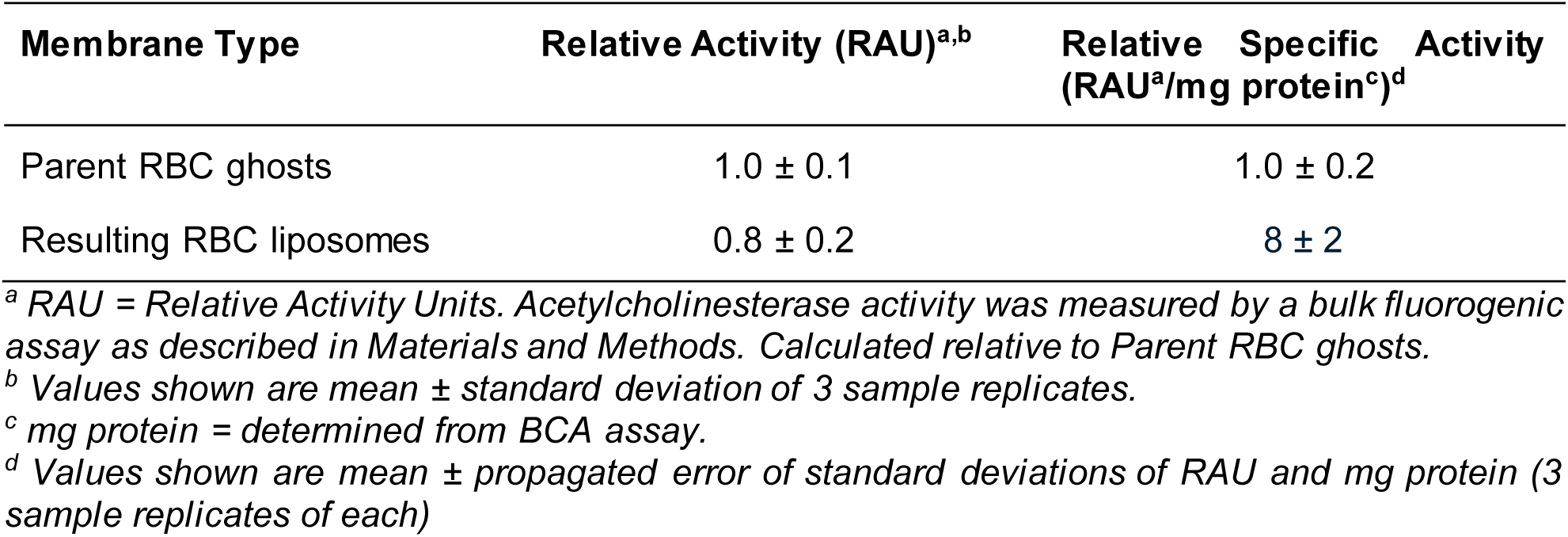
Relative activity and specific activity of acetylcholinesterase in RBC ghosts and RBC liposomes.

In a separate assessment of the functionality of the RBC-SLBs, we quantified the binding of Sendai virus (SeV) to RBC-SLBs compared to control SLBs composed of rupture vesicles only (**Figure 6**). Sendai virus is a well-studied member of the paramyxovirus family, causing disease in rodents and other animals ^28^, and is the subject of ongoing work in our laboratory ^29,30^. Sendai virus is known to bind to red blood cells, utilizing sialic acid glycolipids and glycoproteins (including glycophorin A, ^31^) as attachment receptors. To quantify binding of Sendai virus to the SLBs, we introduced fluorescently labeled Sendai virus particles into our microfluidic flow cells, and quantified the binding of individual virions by fluorescence microscopy after 10 minutes, using an assay we have described previously ^29^. We observed substantial binding of the Sendai virus particles to our RBC-SLBs, with minimal binding to our control SLBs (**Figure 6**). These results indicate that our RBC-SLBs can be used as functional targets for binding experiments.

**Figure 6.**
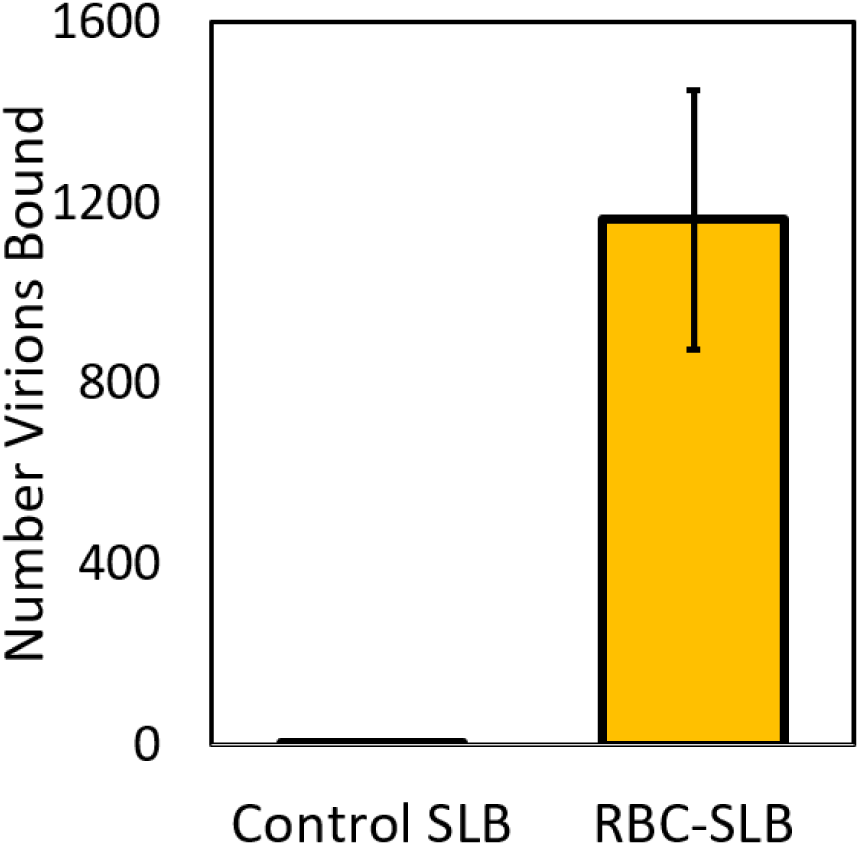
Sendai virus binding demonstrates functionality of RBC-SLBs. Fluorescently-labeled Sendai virus was introduced into a microfluidic device following formation of either a RBC-SLB formed using the rupture vesicle approach, or a control SLB form from rupture vesicles only. Shown are the number of bound virions in a microscope field-of-view after 10 minutes. Values shown are average ± standard deviation of 3 sample replicates, with ≥10 image locations in each sample.

### Preparation of RBC tethered liposomes

Our second model lipid membrane prepared from RBCs was tethered liposomes. To prepare tethered liposomes, we utilized a biotin-NeutrAvidin tethering approach on a PLL-PEG polymer layer (**Figure 7A**). First, we prepared labeled RBC liposomes with biotinylated lipids in addition to the Oregon Green fluorescent label (see schematic in **Figure 1**). Separately, we treated a glass coverslip with PLL-PEG/PLL-PEG-biotin polymer; this results in the PEG/PEG-biotin portion of the co-block polymer being exposed to the aqueous solution, with the PLL portion of the co-block adhered electrostatically to the negatively charged glass coverslip. We then introduced a solution of NeutrAvidin, which bound to the exposed biotin. Finally, we injected the labeled RBC liposomes into the flow cell, which bound to the opposite face of the NeutrAvidin protein, thereby tethering the liposomes. We could then visualize these tethered liposomes by fluorescence microscopy (**Figure 7B**), observing isolated punctate spots whose density could be controlled by altering either the concentration of the RBC liposomes, or by injecting multiple rounds of the RBC liposomes at a set concentration.

**Figure 7.**
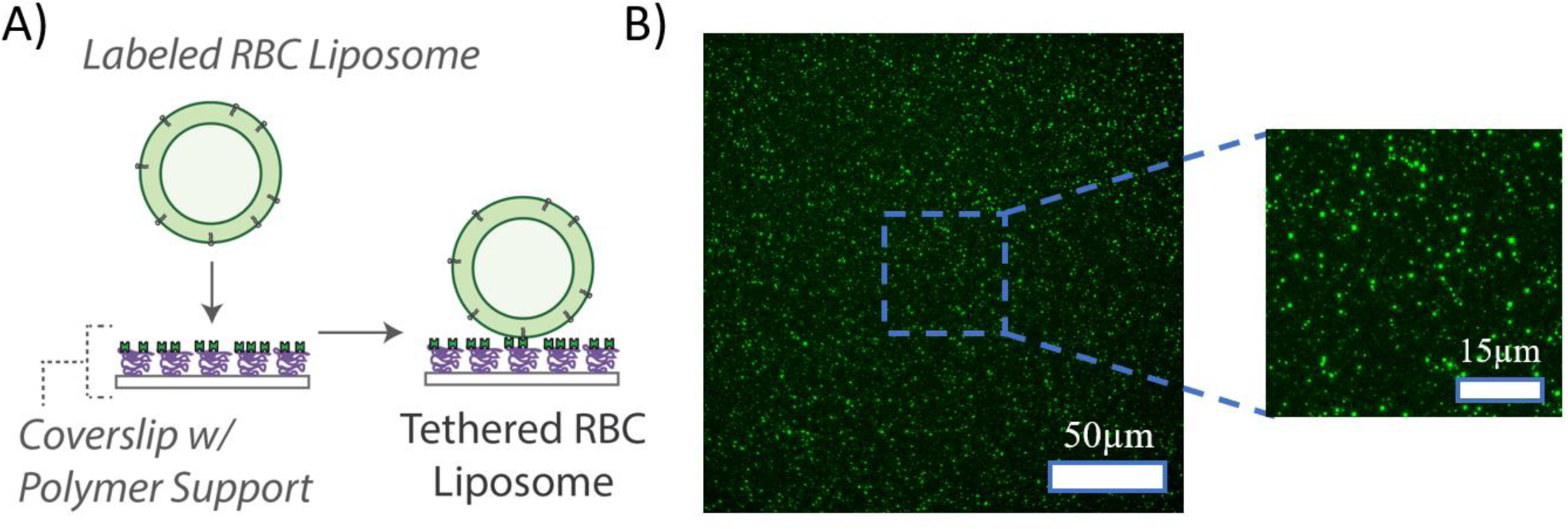
Preparation of tethered RBC liposomes. A) Schematic of preparation of tethered RBC liposomes. RBC liposomes labeled with both Oregon Green-DHPE and biotin-DPPE lipid are added to a polymer-coated glass coverslip inside a microfluidic device. The polymer support is composed of PLL-PEG doped with PLL-PEG-biotin to which NeutrAvidin has been bound. Labeled RBC liposomes then bind to the exposed NeutrAvidin. Unbound liposomes are removed by buffer rinsing. B) Example fluorescence micrograph (with zoomed inset) of tethered RBC liposomes.

### Characterization/validation of RBC tethered liposomes

We conducted a number of different experiments to validate and characterize the RBC tethered liposomes. Some experiments were similar to those conducted for RBC-SLBs, as described above.

First, to verify proper tethering of the RBC liposomes, we compared the number of tethered liposomes bound when the RBC liposomes were prepared with and without biotinylated lipid (**Figure 8**). We observed a large ∼10x increase in the number of tethered liposomes when the RBC liposomes were prepared with biotinylated lipid, indicating that incorporation of the biotinylated lipid and proper tethering had taken place.

**Figure 8.**
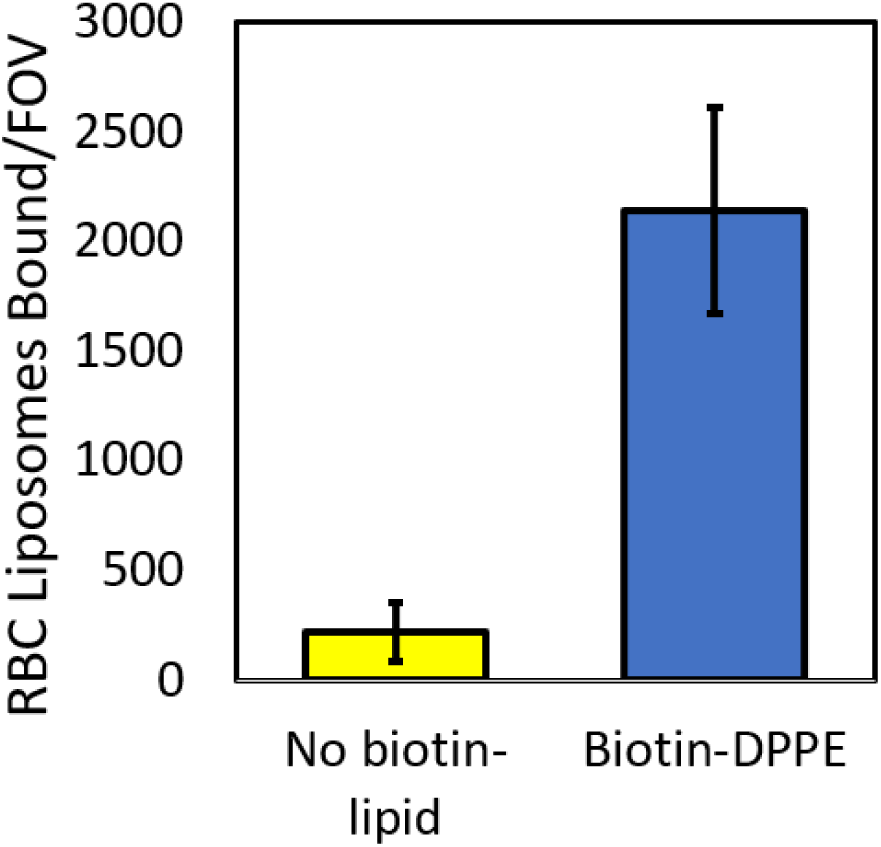
Tethering of RBC liposomes requires incorporation of biotin-lipid. Oregon Green-labeled RBC liposomes were prepared with and without biotin-DPPE lipid, and then tethered to a polymer supported coverslip as described (see Figure 7A). Shown are the number of tethered RBC liposomes per microscope field-of-view (FOV), mean ± standard deviation of ≥20 image areas across 2 sample replicates.

**Figure 9.**
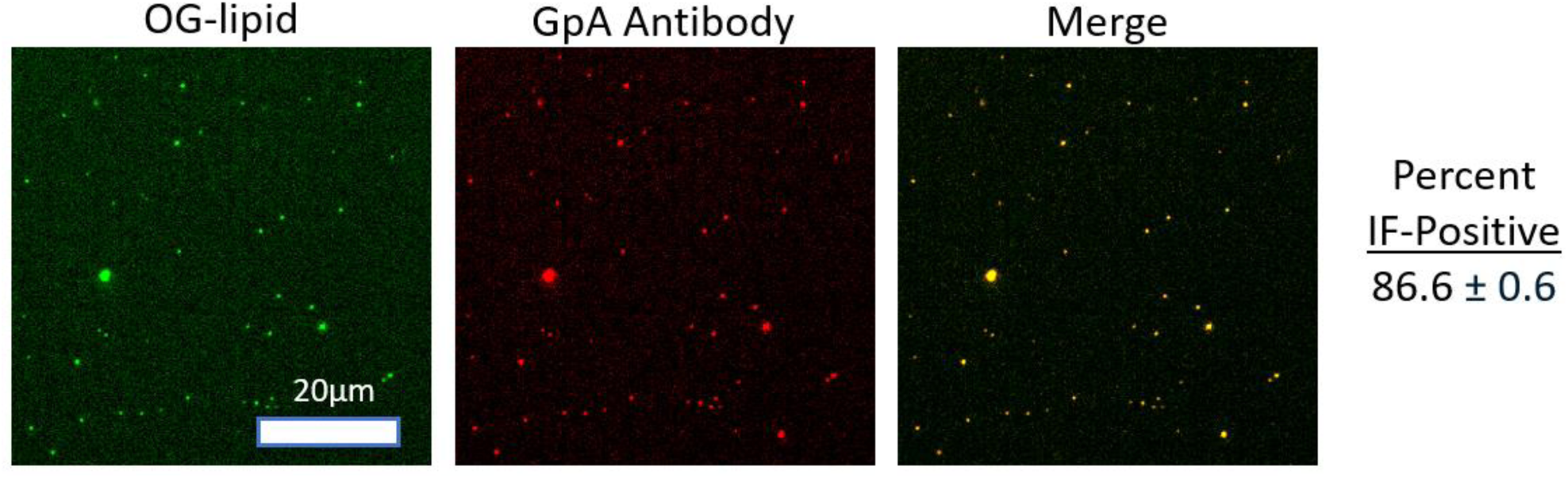
Tethered RBC liposomes exhibit a high degree of IF-positivity for GpA. Tethered RBC liposomes were immunofluorescently (IF) labeled for the glycophorin A (GpA) transmembrane protein (Primary antibody = mouse IgG anti -glycophorin A (CD235a), secondary antibody = goat anti-mouse IgG with Alexa 647). Shown are example images in the Oregon Green-lipid channel (RBC membrane dye), the Alexa 647 channel (GpA Antibody), and merged. The percent IF-positive was calculated as the percentage of RBC liposomes in the green channel co-localized with GpA antibody in the red channel. The value shown is the average ± standard deviation of 3 sample replicates, with ≥8 image locations in each sample.

As an important aside, we note that a key step in the tethering procedure is the proper incorporation of the biotin-lipid into the RBC ghosts. At high concentrations (≥1 g/L), the biotin-lipid is dissolved in a solvent mixture composed of 65/35/8 chloroform/methanol/water (v/v/v), as recommended by the manufacturer. However, at low concentrations (∼0.25 g/L) it can be diluted into ethanol. We found that dilution into ethanol was essential to ensure proper incorporation of the biotin-lipid into the RBC ghosts. Attempts to incorporate the lipid while dissolved in the original solvent mixture proved unsuccessful – little tethering of RBC liposomes was observed in this case (**Figure S4**).

In a second validation experiment, we used immunofluorescence to quantify the fraction of tethered liposomes which properly displayed the glycophorin A membrane protein. We observed that nearly all (∼90%) of the tethered liposomes were IF-positive, indicating the presence of glycophorin A in all or nearly all RBC liposomes. This matches our expectations from a back of the envelope calculation, as follows. The surface area of a human red blood cell is reported at ∼140 µm² ^32^ and the number of glycophorin A proteins per RBC ∼1 x 10^6^ ^26^. The surface area of a ∼200 nm diameter RBC liposome (the average diameter determined by DLS, see **Figure S1**) is calculated as 4*πr*^2^ = 0.13 µm², which means that the number of RBC liposomes produced per RBC is approximately 140/0.13 = 1.1 x 10^3^. Therefore, assuming an even distribution of glycophorin A among the resulting RBC liposomes, the average number of glycophorin A per liposome is estimated as (1 x 10^6^)/(1.1 x 10^3^) = 910. So, even with statistical fluctuations, we might expect all liposomes to have many copies of glycophorin A, and therefore to be IF-positive under ideal assay conditions. This is consistent with our observations.

Finally, we assessed the functionality of the tethered RBC liposomes using the acetylcholinesterase assay described above, comparing to control liposomes composed solely of synthetic lipids (68.95/20/10/1/0.05 POPC/DOPE/Chol/Biotin-PE/OG-DHPE) with no AchE enzyme (**Figure 10**). As with the RBC-SLBs, we observed a substantial increase in the fluorescent product for the tethered RBC liposomes compared to the control, indicating robust activity of the acetylcholinesterase enzyme in the tethered RBC liposomes. Additionally, as discussed above with the RBC-SLBs, while there is no obvious positive control sample to standardize the activity levels, we note that precursor RBC liposomes possessed a similar level of acetylcholinesterase activity to the labeled RBC ghosts, as discussed previously (see **Table 1**).

**Figure 10.**
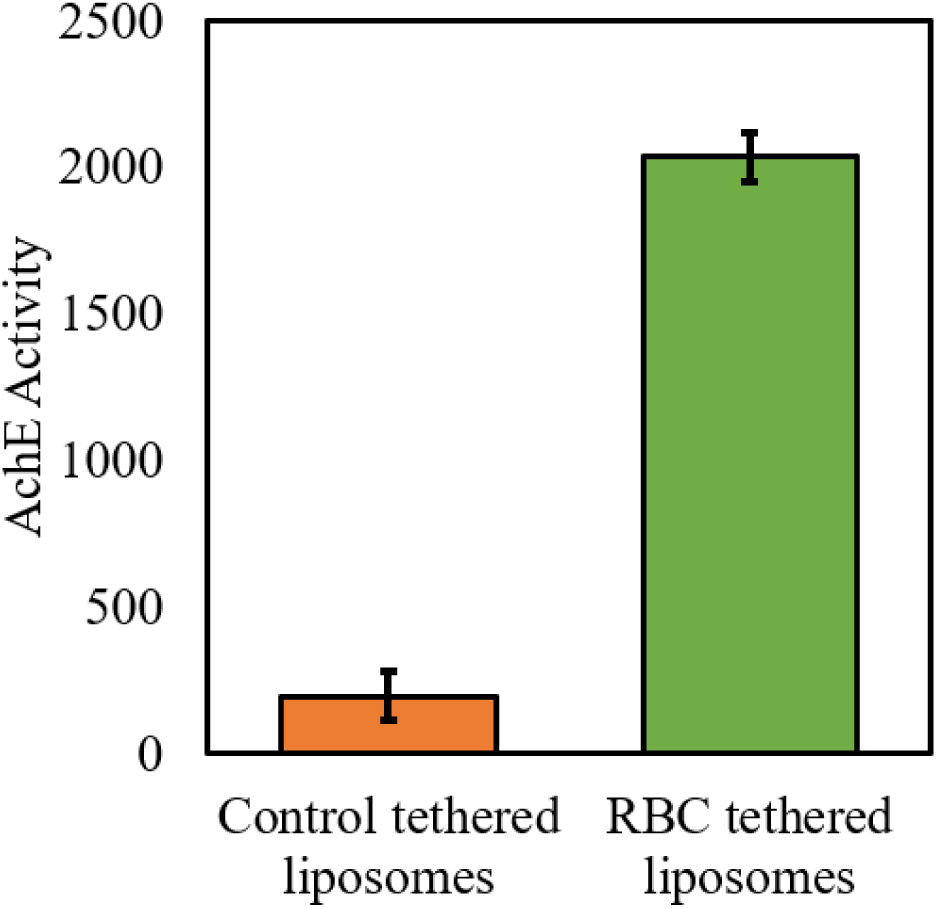
Acetylcholinesterase (AchE) activity of tethered RBC liposomes demonstrates that membrane-anchored enzymes are active. The enzymatic activity of tethered RBC liposomes was assessed by a microscope-adapted red fluorogenic AchE assay (see Materials and Methods for details). This was compared to the activity of control tethered liposomes – synthetic liposomes with no AchE (lipid composition = 68.95% POPC, 20% DOPE, 10% Chol, 1% biotin-DPPE, 0.05% OG-DHPE). AchE Activity is the average red fluorescence within a microscope field-of-view near the tethered liposome surface after 5 minutes at RT. Values shown are average ± standard deviation of 3 sample replicates, with ≥10 image locations in each sample.

### Preparation of Tethered RBC Ghosts

Finally, we briefly explored an alternative model membrane platform – tethered RBC ghosts – that is a variation of the tethered RBC liposomes. Tethered RBC ghosts were prepared in the same manner as was done for the tethered liposomes described above (**Figure 1**), but with the RBC ghosts themselves being tethered onto the PLL-PEG-biotin coated coverslip (**Figure 11**), rather than first being extruded into liposomes. We observed, however, that an important modification was needed in order to successfully create this membrane platform: a biotinylated lipid with a PEG(2000) linker was required. When we incorporated a biotin-lipid with no linker (biotin-DPPE) into the RBC ghosts, the ghosts were observed to loosely attach to the surface, but were not stable upon rinsing (**Figure 11B**). If however we incorporated biotin-PEG(2000)-DSPE into the RBC ghosts, stable binding could be achieved. Presumably this occurred due to the longer linker enabling better accessibility of the biotin to the NeutrAvidin, which in turn increased the number of tethers to a high enough density such that stable binding could be achieved. Tethered RBC liposomes on the other hand likely required many fewer tethers in order to be stably bound such that the biotin-lipid with no linker was sufficient in that case.

**Figure 11.**
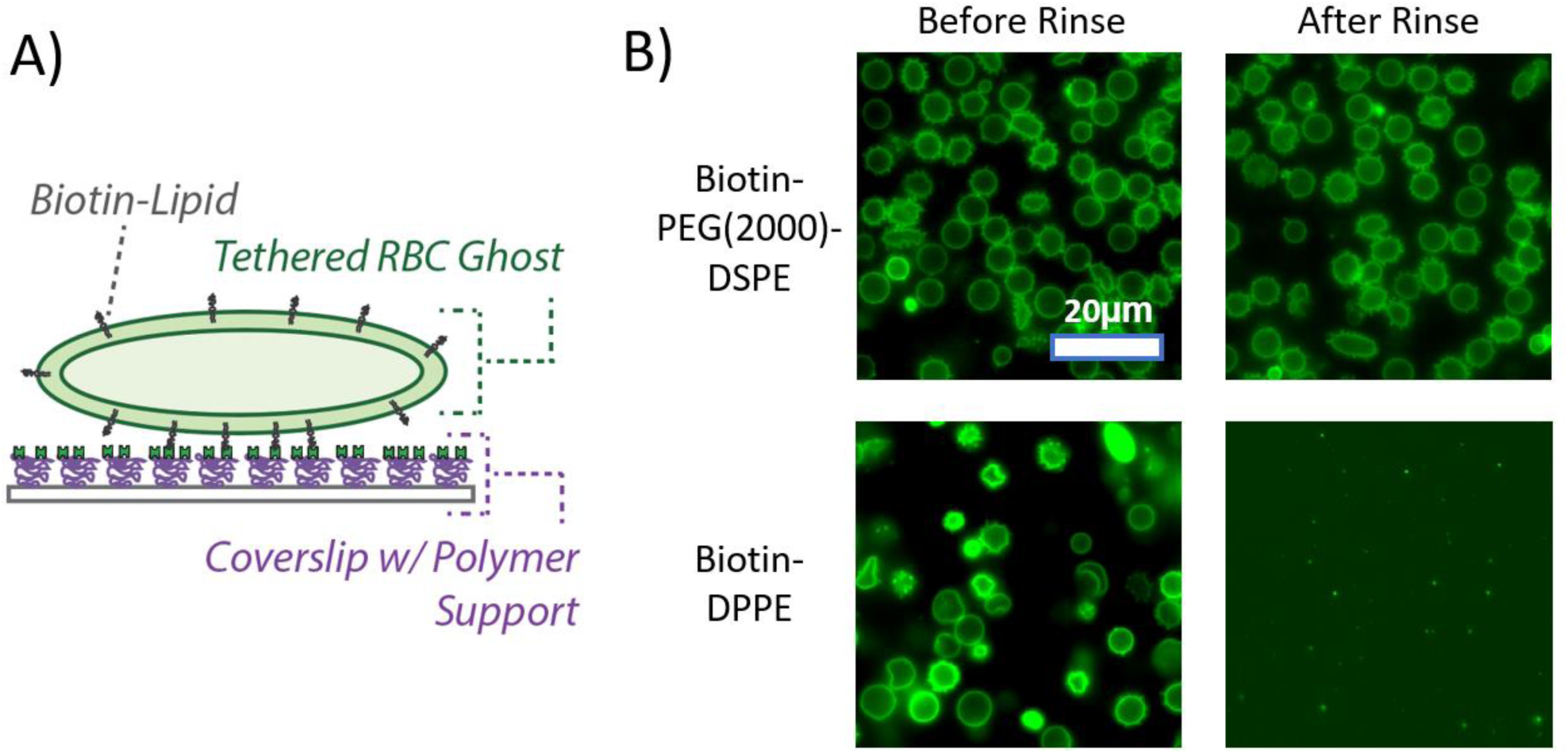
Preparation of tethered RBC ghosts. A) Schematic of preparation of tethered RBC ghosts. RBC ghosts labeled with both Oregon Green-DHPE and biotin-lipid are added to a polymer-coated glass coverslip inside a microfluidic device. The polymer support is composed of PLL-PEG doped with PLL-PEG-biotin to which NeutrAvidin has been bound. Labeled RBC ghosts then bind to the exposed NeutrAvidin. Unbound or loosely bound ghosts are removed by buffer rinsing. B) Example fluorescence micrographs of tethered RBC ghosts before and after rinsing. Tethered ghosts were prepared from ghosts which had been labeled either with biotin-PEG(2000)-lipid or biotin-DPPE (no PEG linker). Only the ghosts labeled with biotin-PEG(2000)-lipid were stable upon rinsing.

## Conclusion

We have presented approaches to prepare two different model membranes derived from human red blood cells – supported lipid bilayers (RBC-SLBs) and tethered RBC liposomes.

For the RBC-SLBs, we found that employing the “rupture vesicle” strategy described by Daniel and coworkers^9,10^ was successful in forming SLBs from RBCs, as demonstrated by lipid mobility assessed by FRAP. The acetylcholinesterase enzyme remained active in the SLB, and the glycophorin A membrane protein was well distributed as observed by immunofluorescence. Additionally, binding experiments with Sendai virus demonstrated that the RBC-SLBs could be used as functional targets for binding. However, increasing the fraction of SLB that was derived from the RBC resulted in a smaller mobile fraction of the lipids, and immunofluorescence FRAP suggested that the glycophorin A was largely immobile. Depending on the given research application, such features may be limitations of this RBC-derived membrane platform.

Tethered RBC liposomes were successfully prepared using a biotin-NeutrAvidin tethering approach after incorporating biotinylated lipids into the RBC liposomes. As with the SLBs, acetylcholinesterase activity persisted in the surface-tethered liposomes, and a high degree of immunofluorescence-positivity for the glycophorin A protein was observed. More briefly, we also demonstrated that tethered RBC ghosts could be prepared in a similar fashion as tethered RBC liposomes, although a PEG linker in the biotinylated lipid was required to achieve stable binding in that case. One advantage of the tethered RBC ghost approach is that it may preserve phospholipid asymmetry in the membrane, as previous work has demonstrated ^33,34^. Such membrane asymmetry may be important for some research applications.

We anticipate that our results and methodologies will be of interest to researchers studying RBCs in a variety of contexts, particularly for researchers interested in studying molecular interactions with RBC membranes.

## Materials and Methods

### Materials

Palmitoyl oleoyl phosphatidylcholine (POPC), dioleoylphosphatidylethanolamine (DOPE), cholesterol (Chol), 1,2-dipalmitoyl-sn-glycero-3-phosphoethanolamine-N-(cap biotinyl) (biotin-DPPE), 1,2-distearoyl-sn-glycero-3-phosphoethanolamine-N-[biotinyl(polyethylene glycol)-2000] (biotin-PEG(2000)-DSPE), 1,2-dipalmitoyl-sn-glycero-3-phosphoethanolamine- N- [methoxy(polyethylene glycol)-5000] (PEG5000-DPPE) and the ganglioside receptor GQ1b were purchased from Avanti Polar Lipids (Alabaster, AL, USA). Poly-L-lysine grafted polyethylene glycol (PLL-g-PEG) and poly-L-lysine grafted biotinylated PEG (PLL-g-PEG-biotin) were obtained from SuSoS AG (Dübendorf, Switzerland). NeutrAvidin protein, Oregon Green 1,2- dihexadecanoyl-sn-glycero-3-phosphoethanolamine (OG-DHPE), and Texas Red 1,2- dihexadecanoyl-sn-glycero-3-phosphoethanolamine (TR-DHPE) were purchased from Thermo Fisher Scientific (Waltham, MA, USA). Chloroform and methanol were acquired from Fisher Scientific (Pittsburgh, PA, USA), while buffer salts and bovine serum albumin (BSA) were sourced from Sigma-Aldrich (St. Louis, MO, USA). 20X phosphate buffered saline was obtained from VWR (Radnor, PA, USA). Polydimethylsiloxane (PDMS) base and curing agent (Sylgard 184) were obtained from Ellsworth Adhesives (Germantown, WI, USA). Anti-glycophorin A primary antibody mouse anti-human CD235a was obtained from Invitrogen (Carlsbad, CA, USA). Goat anti-mouse IgG conjugated to Alexa Fluor 647 and the acetylcholinesterase assay kit (No. ab138871) were obtained from Abcam (Eugene, OR, USA). Human red blood cells (type O+) were obtained from a single donor through Innovative Research (Novi, MI, USA). The cells had been washed with saline to remove the buffy coat and residual debris by the manufacturer.

### Buffer Definitions

● 1X PBS = 137 mM NaCl, 2.7 mM KCl, 10 mM phosphate buffer, pH 7.5

○ Dilutions of 1X PBS to 0.5X, 0.25X, and 0.125X were made using ultrapure water.
● 1X PBSM = 137 mM NaCl, 2.7 mM KCl, 10 mM phosphate buffer, 0.2 mM MgCl₂, pH 7.5
● Reaction buffer (RB pH 7.4) = 10 mM NaH2PO4, 90 mM sodium citrate, 150 mM NaCl, pH 7.4.
● HEPES Buffer (HB pH 7.2) = 20 mM HEPES, 150 mM NaCl, pH 7.2.

All buffers were sterile-filtered at 0.22 μm pore size.

### Fluorescence Microscopy

Fluorescence microscopy images were acquired using Zeiss Axio Observer 7 inverted microscopes (Carl Zeiss Microscopy, LLC., White Plains, NY) equipped with a 63x oil immersion objective (NA = 1.4) and illuminated with a Lumencor Spectra III LED Light Engine or a Lumencor Sola Light Engine. Image acquisition was performed using a Hamamatsu ORCA-Flash 4.0 digital CMOS camera (Hamamatsu Photonics K.K., Hamamatsu City, Japan) set to 16-bit mode, and controlled via Micro-Manager software ^35^. Images and video micrographs were captured at 100 ms per frame with 2×2 pixel binning. For Oregon Green images, filter cube settings were: ex= 475/50 nm, bs = 506 nm, em = 540/50 nm, with a typical light engine intensity setting of 2/1000 (cyan LED). For Texas Red and AbRed images, filter cube settings were: ex = 562/40 nm, bs = 593 nm, em = 641/75 nm, with a typical light engine intensity setting of 10/1000 (green LED). For Alexa Fluor 647 images, filter cube settings were: ex = 640/30 nm, bs = 660 nm, em = 690/50 nm, with a typical light engine intensity setting of 1/1000 (red LED).

### Red blood cell (RBC) ghost preparation

Successive rounds of hypotonic treatment and purification by centrifugation (Sorvall Legend XTR, Thermo Scientific) was used to prepare RBC ghosts. To begin, 400 µL of RBCs were diluted to 10 mL with 0.5X PBS in a 15 mL centrifuge tube, and then pelleted at 4816 × g for 20 minutes at 4°C. A red-tinged pellet was visible after this first spin, indicating a large quantity of hemoglobin was still present. The pellet was then resuspended and washed with 0.25X PBS, followed by repeated washes with 0.125X PBS until pellet appeared pale yellow to white, centrifuging each time for 4816 × g for 20 minutes at 4°C. After the final centrifugation, the supernatant was carefully removed and the pellet was resuspended in 240 µL of 1X PBS.

### Dye and/or biotin labeling of RBC ghosts

For dye labeling, lipophilic OG-DHPE dye (0.075 g/L in ethanol) was added to the resuspended RBC ghosts at a ratio of 3 µL dye per 100 µL of RBC ghosts. For biotin labeling without PEG linker, biotin-DPPE (0.25 g/L in ethanol) was added at a ratio of 2.5 µL biotin-lipid per 100 µL of RBC ghosts. For biotin labeling with PEG linker, biotin-PEG(2000)-DSPE (0.75 g/L in ethanol) was added at a ratio of 2.5 µL biotin-lipid per 100 µL of RBC ghosts. In all cases, the sample was then incubated at room temperature for 2.5 hours in the dark. To remove unincorporated dye and/or biotin-lipid, 1X PBSM was added to the 240uL of labeled ghosts to a final volume of 12 mL and then centrifuged at 4816 × g for 20 minutes at 4°C. The pellet was then resuspended in 240 µL of 1X PBSM.

As described in the Results and Discussion, it is important that the biotin-lipids be diluted into ethanol for proper incorporation into RBC ghosts. At high concentrations (≥1 g/L), biotin-DPPE is typically dissolved in a solvent mixture composed of 65/35/8 chloroform/methanol/water (v/v/v), as recommended by the manufacturer. However, at lower concentrations (∼0.25 g/L) it can be diluted into ethanol.

### RBC liposome preparation

RBC liposomes were prepared by extrusion of RBC ghosts using a mini-extruder (Avanti Polar Lipids, Alabaster, AL). RBC ghosts were first extruded through a 400 nm-pore membrane to remove residual debris left over from the ghost preparation process. If membrane rupture occurred due to a buildup of residual debris, the RBC ghosts were re-extruded through a new 400 nm-pore membrane. These were then extruded through a 100 nm-pore membrane 21 times. RBC liposomes were stored at 4°C and used within approximately one week.

### Preparation of synthetic vesicles (including rupture vesicles)

Synthetic vesicles were prepared using thin-film hydration extrusion, as previously described ^29^. Briefly, purified lipids in chloroform/methanol were added to a test tube with a glass syringe at the desired molar ratio and dried to a film using N2 gas, followed by house vacuum for a minimum of 2 hours. For rupture vesicles, the standard lipid composition was 99.5 mol% POPC and 0.5% PEG5000-DPPE (3.2 x 10^-7^ moles of total lipid). The lipid compositions for other synthetic vesicles (typically 1.4 x 10^-7^ moles of total lipid) are noted together with their respective data. 250 μL of 1.0X PBSM was added to the dried lipid film and rehydrated for approximately ten minutes. The hydrated film was then vortexed at maximum speed for at least sixty seconds to ensure complete lipid recovery ^36^. The resulting suspension was extruded 21 times through a mini-extruder with a 50 nm-pore membrane (for rupture vesicles used for SLB prep) or 100 nm-pore membrane (vesicle used for tethered vesicles). The resulting vesicles were then stored at 4°C and used within approximately one week.

### Microfluidic device preparation

PDMS microfluidic devices affixed to a glass coverslip were prepared as previously described ^29^. This process requires 3 steps: preparation of PDMS flow cells, cleaning of glass coverslips, and plasma-activated bonding.

1. **Preparation of PDMS flow cells:** Briefly, flow cell molds were prepared by affixing small strips (dimensions = 2.5 mm x 13 mm x 70 µm) of Kapton polyimide tape (Ted Pella Inc., Redding, CA) to a glass microscope slide inside a petri dish. PDMS was prepared by mixing elastomer base and curing agent in a 10:1 (w/w) ratio, degassing in house vacuum for ∼1 hour, and then pouring over the mold to ∼0.5 in thickness. The PDMS was then cured for 2 hours at 70°C until solid. Flow cells were cut out using a scalpel, and a biopsy hole puncher (2.5 mm diameter, Harris Uni-core, Ted Pella Inc.) was used to create inlet/outlet holes. Total flow cell volume was ∼4 μL. Flow cells were stored at room temperature covered with a small piece of scotch tape to protect from dust.
2. **Glass coverslip cleaning:** Glass coverslips (24 x 40 mm, No. 1.5 VWR International, Randor, PA) were cleaned in a 1:7 solution of 7x detergent (MP Biomedicals, Burlingame, CA) and deionized water, with heating until the solution became clear. The coverslips were then extensively rinsed under running DI water for >2 hours, given a final brief (several seconds) rinse with Millipore water, and then baked at 400°C for 4 hours in a kiln. They were stored at room temperature.
3. **Plasma-activated bonding:** Individual PDMS flow cells (channel side upward) and clean glass coverslips were exposed to air plasma for 60 seconds using a Harrick Plasma Cleaner (Model PDC-3xG, Harrick Plasma, Ithaca, NY). Then, the PDMS chip was immediately inverted and placed channel-side down onto the glass coverslip. This bonding forms a leak-resistant seal between the PDMS and the glass. To create a buffer reservoir, the tip of a 1 mL plastic pipette was cut and secured to the designated inlet hole of each microchannel using five-minute epoxy (Devcon, ITW Polymer Adhesives North America, Danvers, MA).

Once prepared, microfluidic devices were immediately used to prepare model membranes.

### RBC-SLB formation

RBC-SLBs were prepared using an adaptation of the rupture vesicle approach, as reported by the Daniel lab ^9,10^. Immediately after microfluidic device assembly, 5 µL of RBC liposomes (typical liposome concentration ∼0.5-5 nM depending on desired density) was immediately added to the flow cell channel by pipette. Additional injections of 5 µL could be added if even higher densities were desired. The RBC suspension was incubated inside the channel for an hour at room temperature. The channel was then rinsed with 1.5 mL of 1X PBSM buffer using a Fusion 200 syringe pump (Chemyx Inc., Stafford, TX, USA), flow rate 800 µL/min. Then, rupture vesicles (total lipid concentration = 0.4 mg/mL) were added, incubated for an hour, and then rinsed with 1.5 mL of 1X PBSM.

### Tethered RBC liposome and tethered RBC ghost preparation

A 95:5 (v/v) solution of PLL-g-PEG (1 g/L in HB pH 7.2) and PLL-g-PEG-biotin (1 g/L in HB pH 7.2) was prepared fresh in an Eppendorf tube. Immediately after microfluidic device assembly, 5 μL of this solution was added into the flow cell channel by pipette, and incubated for 30 minutes at room temperature. The channel was then rinsed sequentially with 1.5 mL of Milli-Q water and 1.5 mL of 1X PBSM buffer by syringe pump, flow rate 800 µL/min. 6 μL of 0.2 mg/mL NeutrAvidin solution in RB pH 7.4 was then added. After a 15-minute incubation, the channel was rinsed again with 1.5 mL of 1X PBSM buffer. 6 μL of liposome or RBC ghost suspension was pulled into the channel. After a 1-hour incubation at room temperature, a final rinse with 1.5 mL of 1X PBSM buffer was performed.

### Fluorescence recovery after photobleaching (FRAP)

Fluorescence recovery after photobleaching (FRAP) was employed to assess mobility and diffusion kinetics within SLBs. After focusing the microscope on the SLB using low intensity illumination, the field aperture was closed to illuminate only a small centralized region. This region was then photobleached using high-intensity illumination (maximum light engine intensity setting) for 5 seconds. The field aperture was quickly opened and the subsequent recovery of fluorescence in the bleached area was monitored at low intensity via time-lapse imaging. The average fluorescence intensity over time within the photobleached region was extracted using FIJI ^37^. The average intensity within a background region far from the photobleached area was also extracted to quantify residual photobleaching that occurs during the time-lapse imaging itself. These intensity data were then processed using a custom-written MATLAB script to estimate the diffusion coefficient and the fraction immobile (source code at https://github.com/rawlelab/SendaiBindingAnalysis). In the script, the normalized fluorescence recovery curve *I_norm_(t)* within the photobleached area (shown as Fraction Recovered in data figures) was calculated as:

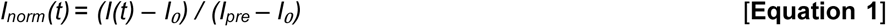

where *I(t)* is the intensity at time *t*, *I₀* is the intensity immediately after photobleaching, and *I_pre_* is the intensity before photobleaching. The script also applied a photobleaching correction by fitting the background ROI signal to a linear decay model. This correction factor was then applied to the *I_norm_(t)*, typically resulting in only very minor changes to the fluorescence recovery curve. This corrected recovery curve was then fit to a theoretical FRAP model based on the classic equation by Soumpasis ^38^:

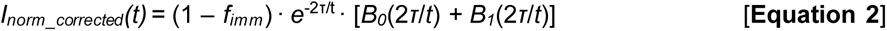

where *f_imm_*is the immobile fraction, *τ* = *r*² / 4*D* is the characteristic recovery time, *r* is the radius of the photobleached spot, *D* is the diffusion coefficient, and *B_0_* and *B_1_*are modified Bessel functions of the first kind (orders 0 and 1).

### Immunofluorescence measurements of RBC model membranes

RBC-SLBs or tethered RBC liposomes were prepared in a microfluidic flow channel as described above. To the flow channel, 4 µL of 30 mg/mL BSA in 1X PBSM was added to prevent nonspecific antibody binding. After a 15 minute incubation at room temperature, 4 µL of primary antibody was pulled through the channel and incubated for 15 minutes. Primary antibody = 1:50 (SLBs) or 1:25 (tethered liposomes) dilution in 1X PBSM of mouse anti-human glycophorin A (CD235a) at 0.5 mg/mL. Then, the channel was rinsed with 2 mL of 1X PBSM by syringe pump, flow rate = 0.8 mL/min. A second blocking step was then performed by pulling through 4 µL of 30 mg/mL BSA and incubating for 15 minutes. 4 µL of secondary antibody was then pulled through the channel and incubated for 15 minutes. Secondary antibody = 1:50 (SLBs) or 1:100 (tethered liposomes) dilution in 1X PBSM of goat anti-mouse IgG-Alexa Fluor 647 (Abcam AB150115) at 2 mg/mL. Finally, the channel was rinsed with 3 mL of 1X PBSM by syringe pump, flow rate = 0.8 mL/min. The sample was then ready for imaging. For image analysis of SLBs, the average intensity within a field of view was quantified using FIJI ^37^. For analysis of tethered liposomes, the percentage of Oregon Green-labeled RBC liposomes colocalized with Alexa 647 antibody signal was determined using custom-written Matlab scripts, which have been previously described ^29^ and are available at https://github.com/rawlelab/SendaiBindingAnalysis. Briefly, the scripts identify bound liposomes in the green fluorescence channel, and then quantify the fluorescence in the same ROI in the Alexa 647 channel. If the intensity in the Alexa 647 channel is above background, then the liposome is counted as IF -positive.

### Microscope-adapted acetylcholinesterase assay

The microscope-adapted acetylcholinesterase (AChE) assay was adapted from a commercial fluorometric kit (Abcam, Cat. No. ab138871) to use with RBC model membranes in our microfluidic flow cell setup. First, RBC-SLBs, tethered RBC liposomes, or the appropriate control membrane, were prepared in a microfluidic flow channel as described above. Separately, AChE assay solution was prepared by combining 0.5 mL of acetylcholinesterase probe solution (Component B, reconstituted in Assay Buffer, Component E), 0.5 µL of AbRed Indicator stock solution (250X in DMSO), and 2 µL of acetylcholine stock solution (1000X, reconstituted in Assay Buffer, Component E). As recommended by the manufacturer, this AChE assay solution was kept on ice or at 4 °C and used within 4 hours. 15 µL of AChE assay solution was added to each flow cell with a RBC model membrane. After 5 minutes, fluorescence imaging (AbRed channel) was performed in the flow cell immediately adjacent to the model membrane surface to detect AChE activity. To quantify the extent of activity, the average intensity within a field of view was quantified using FIJI (12).

### Bulk acetylcholinesterase assay and BCA measurement

The bulk acetylcholinesterase assay was used to quantify the relative activity of RBC liposomes and parent RBC ghosts. It was performed using a colorimetric fluorometric kit (Abcam, Cat. No. ab138871) according to the manufacturer’s instructions, and measured using a BioTek SynergyHT microplate reader (Agilent Technologies, Winooski, VT, USA), with ex = 530/25 nm, em = 590/35 nm. A commercial BCA assay kit (Pierce Micro BCA Protein Assay Kit) was used to quantify the total protein concentration of RBC liposomes and parent RBC ghosts. It was performed according to manufacturer’s instructions, and quantified using a NanoDrop 2000C (Thermo Scientific, Waltham, MA, USA). Parent RBC ghosts refers to an aliquot of RBC ghosts taken from the same stock that was used to prepare the RBC liposomes.

### Sendai virus binding assay

Single virus binding measurements to SLBs were performed as previously described ^29^. Briefly, a RBC-SLB was prepared in a microfluidic flow channel as described above. Then, 5 µL of Texas Red-labeled Sendai virus was injected into the channel and incubated for 10 minutes at room temperature. Unbound virions were removed by rinsing (1 mL 1X PBSM via syringe pump, flow rate = 0.8 mL/min). Fluorescence images of virions bound to the SLB were collected throughout the flow cell. The number of viral spots in each image was quantified using custom Matlab scripts ^29^, and available at https://github.com/rawlelab/SendaiBindingAnalysis.

### Estimation of labeled RBC liposome concentration

The concentration of labeled RBC liposomes was estimated using a so-called “splat assay” procedure we have previously described to estimate the concentration of fluorescently-labeled viral particles ^29^. Briefly, Oregon Green-labeled RBC liposomes were diluted to an appropriate concentration (typically 1:500) and mixed 1:1 with Texas Red-labeled synthetic vesicles at a nominally known concentration (synthetic vesicle composition = 0.05% Texas Red-DHPE, 69.95% POPC, 30% chol). This nominally known particle concentration was estimated by the moles of total lipid used to prepare the synthetic vesicles, any dilutions that occurred, and estimates of both the cross-sectional area of the lipids and the average surface area of a single liposome. Importantly, both the RBC liposomes and synthetic vesicles were diluted to very low concentrations such that SLB formation would not occur; instead, individual liposomes would be observed. The mixture of diluted RBC and synthetic vesicles was added to a freshly prepared, empty microfluidic flow channel, and incubated for 10 minutes. During this time, nearly all liposomes adhered non-specifically to the clean glass surface. Images were then taken of RBC liposomes and synthetic vesicles in their respective fluorescence channels. The ratio of bound RBC liposomes to bound synthetic vesicles within a field of view could then be used to estimate the concentration of the RBC liposomes relative to the nominally known synthetic vesicle concentration. This estimation method yielded a particle concentration of ∼5 nM for typical preparations of labeled RBC liposomes. However, we did observe some variability between preps, and so for rigorous applications researchers are encouraged to quantify the concentration of each prep individually, rather than assuming all preps are identical.

## Supporting Information

Additional supporting figures, including DLS measurements of RBC liposomes, FRAP results for SLBs formed solely from RBC liposomes, comparison of estimated diffusion coefficients versus Fraction RBC, and influence of solvent on incorporation of biotin-lipid into RBC liposomes.

## Author Contributions

SM and MM designed experiments, collected data, analyzed data and helped write the manuscript. RJR designed experiments, analyzed data, acquired project funding, provided supervision, and wrote the manuscript. All authors reviewed the manuscript.

## Acknowledgements

The authors thank Amy Lam and Abraham Park (Williams College, MA) for their preliminary work to develop RBC-derived model membranes. The authors gratefully acknowledge financial support from Williams College and NIH grant R15AI171754.

## Declaration of interests

The authors declare no competing interests.

## Notes

### Competing Interest Statement

The authors have declared no competing interest.

