## Supporting Information for "Surface-attached model lipid membranes derived from human red blood cells"

**Co-first author contribution

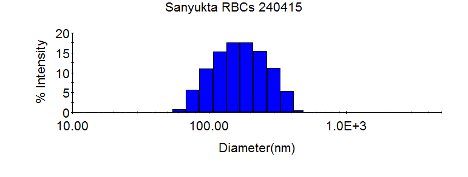

***Figure S1. Dynamic light scattering characterization of labeled RBC liposomes.*** *Mean diameter = 198 nm.*

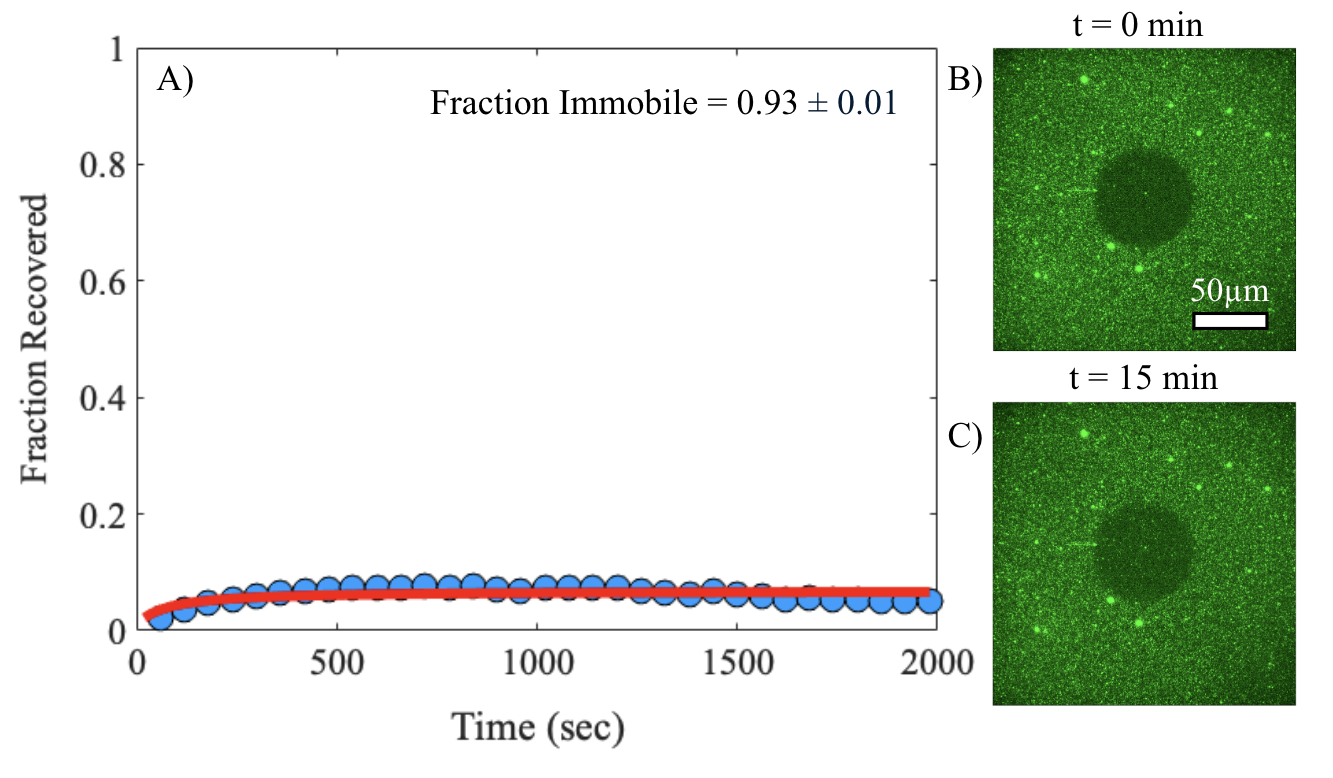

***Figure S2. SLBs formed from RBC liposomes alone exhibit lack of lipid mobility.*** *RBC liposomes labeled with Oregon Green-DHPE were deposited on a glass coverslip to form an SLB. Lipid mobility was assessed by FRAP of the OG-DHPE. A) Example FRAP recovery curve (blue circles = data, red line = fit to FRAP diffusion model,* ***Equation 2*** *in the main text). Fraction recovered is the normalized fluorescence intensity within the photobleached spot, background corrected for residual photobleaching that occurred during the time-lapse imaging. Fraction recovered = 1 was set to the fluorescence intensity immediately prior to photobleaching. Fraction recovered = 0 was set to the fluorescence intensity immediately after photobleaching (t = 0). The immobile fraction calculated from the model fit is shown as the average ± standard deviation of 5 sample replicates. B) and C) Example fluorescence micrographs at t = 0 and t = 15 min, respectively.*

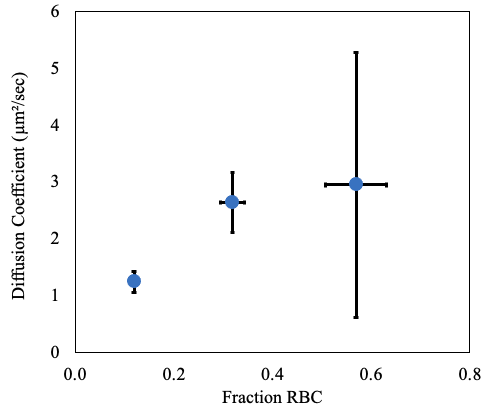

***Figure S3. Plot of estimated diffusion coefficients versus the fraction of SLB composed of the RBC liposomes (Fraction RBC)****. Diffusion coefficients were determined by FRAP diffusion model fits (values shown are average ± standard deviation of 3 sample replicates). Fraction RBC was determined by total image fluorescence comparisons to a standard SLB composed only of rupture vesicles (see main text and Materials and Methods for details). Values shown are average ± propagated error of standard deviations of experimental and standard SLB samples. Standard deviations were calculated from ≥ 20 image locations across 2 sample replicates. Note that the large error vertical error bar at fraction RBC ~ 0.6 is due to the SLB possessing a high immobile fraction (compare to main text* ***Figure 3D****). Hence, the diffusion coefficient estimation is less precise.*

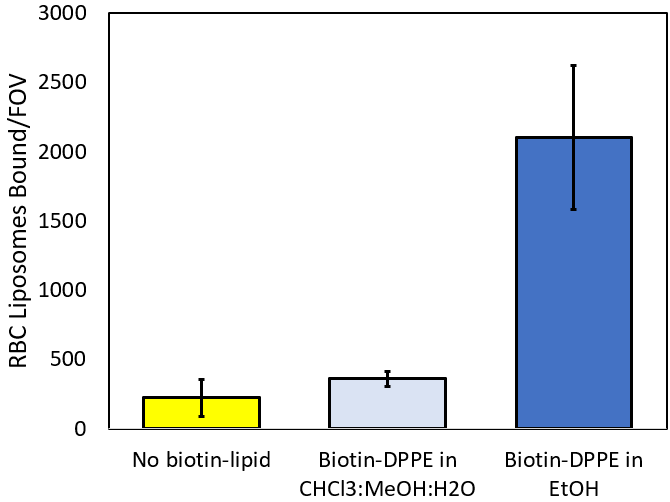

***Figure S4. Proper tethering of RBC liposomes requires incorporation of biotin-lipid dissolved in the proper solvent.*** *Oregon Green-labeled RBC liposomes were prepared either without biotin-DPPE lipid or with biotin-DPPE diluted into two different solvent mixtures – 65/35/8 chloroform/methanol/water (v/v/v, CHCl3:MeOH:H2O) or ethanol (EtOH). The labeled RBC liposomes were then tethered to a polymer supported coverslip as described (see schematic in* ***Figure 7A****). Shown are the number of tethered RBC liposomes per microscope field-of-view (FOV), mean ± standard deviation of ≥8 image areas.*
